# Pre-FIB Layer-Mapping Cryo Tomography (PLCT) for Depth-Resolved in Situ Structural Analysis of Multilayered Tissues

**DOI:** 10.64898/2026.08.25.746966

**Authors:** Fenglan Wang, Xin Lin, Bilin Rao, Xiaoqian Lai, Lihan Yu, Fei Sun, Jia Qu, Jun Zhang

## Abstract

Cryo-electron tomography (cryo-ET) enables near-native visualization of subcellular architectures, yet applying it to moderately thick, multilayered tissues such as the retina is hampered by inadequate vitrification and inaccurate depth-targeting. Here, we developed PLCT, an integrated approach combining modified high-pressure freezing, cryo-ultramicrotome trimming, and plasma-based cryo-FIB milling to overcome these barriers. PLCT reliably vitrified <100 µm retinal strips with minimal ice artifacts, navigates precisely to the outer plexiform layer using morphological landmarks, and produces high-quality lamellae suitable for high-resolution cryo-ET. Subtomogram averaging (STA) analysis identified microtubules at 16.33 Å within retinal horizontal cell processes. Importantly, STA also resolved a 10-nm-diameter filamentous structure at 24.81 Å in the same processes, featuring six peripheral strands surrounding an elongated central density with continuous intervening cavities, an architecture consistent with intermediate filaments. Together with its native localization and immunoreactivity, these features collectively identify the filaments as neurofilaments. Separately, 3D reconstruction of synaptic ribbons uncovered a previously unrecognized “mahjong tile”-like fine ultrastructure. These results demonstrate that PLCT-produced lamellae are of sufficient quality to support structural analysis in native tissue. Although demonstrated on retinal photoreceptor synapses as a proof-of-principle, PLCT is inherently generalizable, with its depth-navigation and vitrification strategies directly applicable to any multilayered tissues. This work establishes PLCT as a robust, reproducible platform for depth-resolved in situ cryo-ET of multilayered tissues.

## Introduction

Cryo-electron tomography (cryo-ET) has emerged as a powerful approach for visualizing three-dimensional subcellular architectures in near-native states, gradually becoming an important bridge connecting molecular structures with cellular functions^1, 2^. By acquiring tilt series and reconstructing tomograms, cryo-ET achieves nanometer-scale resolution, revealing the in situ organization of protein complexes, organelles, and membrane structures without the artifacts introduced by chemical fixation, dehydration, or staining^3–5^. Currently, most cryo-ET applications are restricted to cultured cells or thin specimens, where vitrification and lamella preparation are relatively straightforward^6–10^. Despite these advances, applying cryo-ET to intact thick tissues, especially those of moderate thickness (>100 µm), poses substantial challenges, including non-uniform ice formation and inaccessible target depths^11, 12^. Adequate vitrification currently relies on either plunge-freezing (effective only up to ∼10 µm) or high-pressure freezing (HPF), which can vitrify samples tens to hundreds of micrometers thick but suffers from low throughput, technical complexity, and challenges in subsequent thinning (e.g., cryo-sectioning introduces surface artifacts, while cryo-lift-out is low-throughput and technically demanding)^13–16^. Moreover, even after successful vitrification, precisely preparing lamellae from defined depths within a thick tissue using cryo-focused ion beam (cryo-FIB) milling remains difficult^17, 18^. Conventional gallium-based FIB lacks sufficient milling efficiency for large volumes and may alter sample architecture, whereas xenon plasma-based FIB, though promising, is still immature in biological contexts^19–21^. Thus, achieving both reliable vitrification of moderately thick tissues and targeted thinning at specific depths represents a major unmet need for in situ tissue cryo-ET.

The retina is the first organ that perceives vision. It is a multilayer tissue composed of ten distinct layers, including three soma layers and two synaptic layers. Within this laminated architecture, the photoreceptor ribbon synapse is situated in the outer plexiform layer (OPL), where it forms the triadic synaptic junction between photoreceptor terminals and the dendrites of bipolar and horizontal cells. As the first synapse in the entire visual system, this ribbon synapse plays an irreplaceable role in transmitting visual signals^22, 23^. However, the mechanism by which its specialized synaptic structure regulates synaptic vesicle release remains unclear. Resolving this complex architecture through in situ cryo-ET may elucidate this mechanism.

To achieve this goal with cryo-ET, the two technical bottlenecks mentioned above, namely, reliable vitrification of moderately thick tissues and targeted thinning at specific depths, must first be overcome. When applying cryo-ET to the retina, these bottlenecks become particularly acute, giving rise to three major difficulties. First, the mouse retina is approximately 200 µm thick, far exceeding the effective range of plunge-freezing. Even with HPF, uniform vitrification across the full thickness is difficult to achieve^24^, with differential ice crystal formation risks between the inner (vitreous-facing) and outer (choroid-facing) layers that can easily disrupt delicate structures such as synapses and membrane systems. Second, the photoreceptor synapses of interest in this study are located at an intermediate depth within the mouse retina, approximately 90–100 µm from either surface. Because this depth is neither the top nor the bottom of the tissue, it cannot be easily accessed by conventional cryo-FIB milling approaches, which typically start from the block surface and mill downward progressively. Without a method to precisely locate and target this specific depth under cryogenic conditions, it is nearly impossible to generate lamellae that contain the photoreceptor synaptic layer. Furthermore, because the retina is highly laminated with two synaptic- and three nuclear-layers, high depth precision is critical; even minor targeting errors can shift the analysis to entirely different structures.

To overcome both general bottlenecks in cryo-ET tissue sample preparation and retina-specific challenges in vitrification and depth-targeted localization, we developed pre-FIB layer-mapping cryo tomography (PLCT), an adaptive and unified approach for targeted cryo-ET of retinal tissues. This method breaks through current technical barriers and provides a reliable basis for in situ structural analysis of all retinal subcellular features, from photoreceptor outer segments to deep-layer synapses and inner plexiform connections. Specifically, rather than attempting to vitrify the full 200 µm thickness of the retina in one piece, we will first dissect the retina into thin strips of less than 100 µm in thickness. These thinner strips are then subjected to HPF, which ensures rapid and uniform vitrification across the entire sample volume, effectively eliminating ice crystal artifacts that commonly plague thicker tissues. Following vitrification, we will employ a cryo-ultramicrotome to perform coarse and fine trimming of the frozen strips, reducing them to approximately 30–50 µm thick sections. This intermediate thinning step bridges the gap between bulk tissue and electron-transparent lamellae, while preserving native structural integrity. Critically, to enable precise targeting of the photoreceptor synapses, which reside at a depth of 90–100 µm within the intact retina, we will first identify the retinal inner and outer sides of the trimmed sample based on morphological landmarks visible under cryo-ultramicrotome. Subsequently, we will transfer the sample to a cryo-FIB/SEM (Hydra, equipped with multimodal imaging capabilities) and further refine the identification to locate the OPL, which contains the photoreceptor synapses. This stepwise depth-navigation strategy ensures that lamellae can be milled exactly at the desired retinal layer, rather than blindly. Once the target layer is confirmed, we will proceed with xenon/argon plasma-based cryo-FIB milling to generate high-quality lamellae suitable for cryo-ET imaging. Upon completion, this method will enable, for the first time, direct three-dimensional visualization and quantitative analysis of native-state synaptic elements within intact retinal tissue—specifically, photoreceptor ribbon synapses in the OPL, and bipolar and amacrine cell synapses in the inner plexiform layer (IPL). Critically, application of this approach has already yielded, also for the first time, in situ reconstruction and analysis of neurofilaments (24.81 Å) and microtubules (16.33 Å) within postsynaptic horizontal cell processes, in addition to successful segmentation of presynaptic photoreceptor synaptic ribbon. Beyond the retina, this approach holds promise for extension to other stratified organs, such as the cerebral cortex, cerebellar cortex, hippocampus, spinal cord, and kidney.

## Result

### Modified HPF yields reliable and unified tissue protection

We performed a careful comparative analysis of three retinal preparation methods for HPF. In the first approach, direct HPF of retinal whole-mounts (∼200 µm thick) resulted in severe ice crystal damage to the ultrastructure and poor preservation of deeper tissue (Supplementary Fig. 1a, a1). In the second approach, which involved agarose embedding followed by vibratome sectioning into <100 µm strips, the prolonged processing increased the risk of tissue degradation. Moreover, agarose infiltration altered tissue morphology and introduced foreign material, compromising the native ultrastructure (Supplementary Fig. 1b, b1). In contrast, only the third (ultimately adopted) approach, rapid dry-cutting of the retina into <100 µm strips (Fig. 1a), achieved minimal processing time, excellent ultrastructure preservation, and negligible ice crystal formation (Supplementary Fig. 1c, c1).

**Figure 1.**
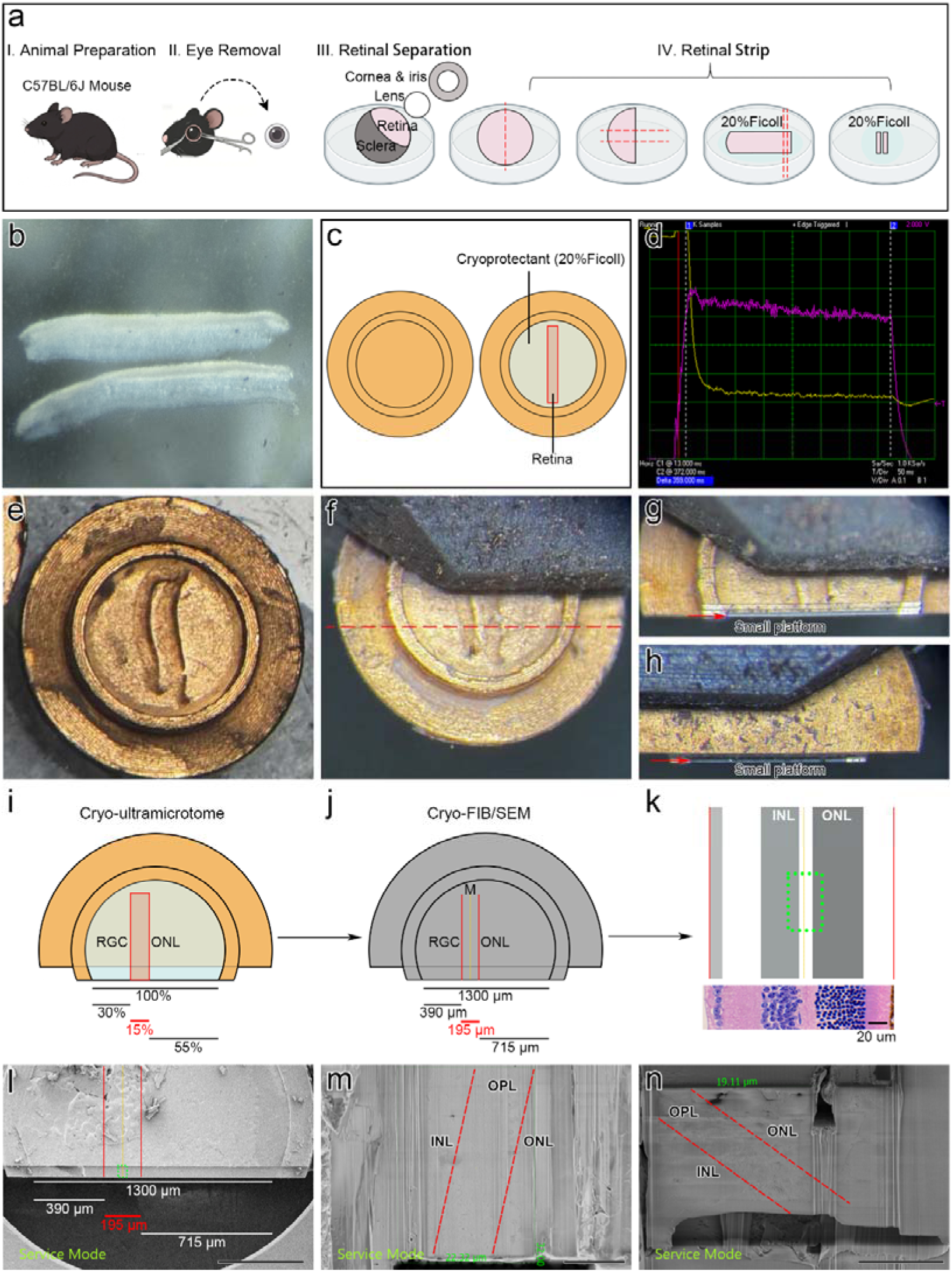
High pressure freezing (HPF), pre-trimming and identifying ROI. **a**, Schematic illustration of the modified tissue acquisition. The retina was rapidly trimmed into ∼100- µm-thick strips with a scalpel. **b**, Representative retinal strips (1.5 mm × 200 µm × 100 µm) are trimmed to fit the HPF carrier dimensions. The schematic depicts a strip centrally positioned within the carrier, and the interstitial space around it is filled with cryoprotectant. **c**, Schematic illustration of a retinal strip centered in the center of the HPF carrier, with cryoprotectant (20% Ficoll) filling the surrounding interstitial space. **d**, Pressure and temperature profiles during HPF, showing controlled vitrification under standard conditions. **e-h**, Cryo-ultramicrotome pretrimming. The sapphire plate and auxiliary ring are removed from the carrier (**e**). The retinal strip is oriented perpendicular to the glass knife edge and the red dashed line indicates the region targeted for rough trimming (**f**). Representative bright-field image of the pretrimmed carrier viewed through the eyepiece, the red arrow indicates the small platform (**g, h**). **i-k**, Schematic illustration of the targeting strategy. The retinal strip position along the carrier x-axis is measured as a percentage using ImageJ (**i**) and then converted to an absolute coordinate in the Hydra system (**j**). The outer plexiform layer (OPL) is identified based on the stratified retinal architecture (**k**). The red line indicates the position of the retinal strip, while the green box denotes the inferred OPL region. **l**, Representative ROI localization in the Hydra system. The red line marks the retinal strip position (converted from cryo-ultramicrotome percentage measurements), and the green box highlights the inferred OPL region selected for subsequent coarse milling. Scale bars: 500 µm. **m**, **n**, Top-view (**m**) and side-view (**n**) of cryo-FIB images after coarse milling and fine polishing show the OPL, located between the outer nuclear layer (ONL) and inner nuclear layer (INL), with the region delineated by the red dashed boundary. Scale bars: 10 µm.

### Percentage-based cryo-ultramicrotome measurement of carrier and retinal tissue

After pre-trimming (see Methods), images were captured through the eyepiece (Fig. 1g, h), and the corresponding distances (percentage) of the retinal strip along the carrier x-axis were measured with ImageJ (Fig. 1i and Supplementary Fig. 4). Briefly, the full width of the carrier is set as 100 %. The width of the rectangular retinal tissue fragment is measured, and its right side is confirmed to be the outer nuclear layer (ONL). The distances from the two edges of the retina to the corresponding sides of the carrier are also recorded as percentages of the total carrier width. This provides a relative spatial map of the retina within the carrier before further processing.

### Precise lift-out of retinal tissue containing the OPL

Following above step, the carrier was transferred into the cryo-FIB/SEM. Using the percentage-based measurements of the carrier and retinal tissue as a reference, we measured the absolute width of the carrier and the actual width of the rectangular retinal tissue in micrometers (Fig. 1j-l), with the right side again identified as the ONL. Additionally, the absolute distances from the retinal edges to both sides of the carrier were recorded. These quantitative, dimension-based data complement the percentage-based information from Fig. 1i, enabling precise geometric localization. The direct comparison between these carrier- and retina-based percentage data further validates their alignment and ensure reliable spatial correspondence.

Furthermore, based on this alignment, and to improve efficiency, the target region was narrowed to the OPL by leveraging the distinct cellular stratification of the retina, reducing the cryo-FIB coarse milling target width to 50 µm (green area in Fig. 1k, l).

Guided by coordinates, the sample top (xy) and lateral (xz) faces were coarsely milled and fine-polished with reduced current. Both planes clearly resolved the OPL (between the ONL and the inner nuclear layer, INL; Fig. 1m, n), confirming precise region of interest (ROI) localization. The ROI was then lifted out using an Easylift micromanipulator (Fig. 2a-c).

**Figure 2.**
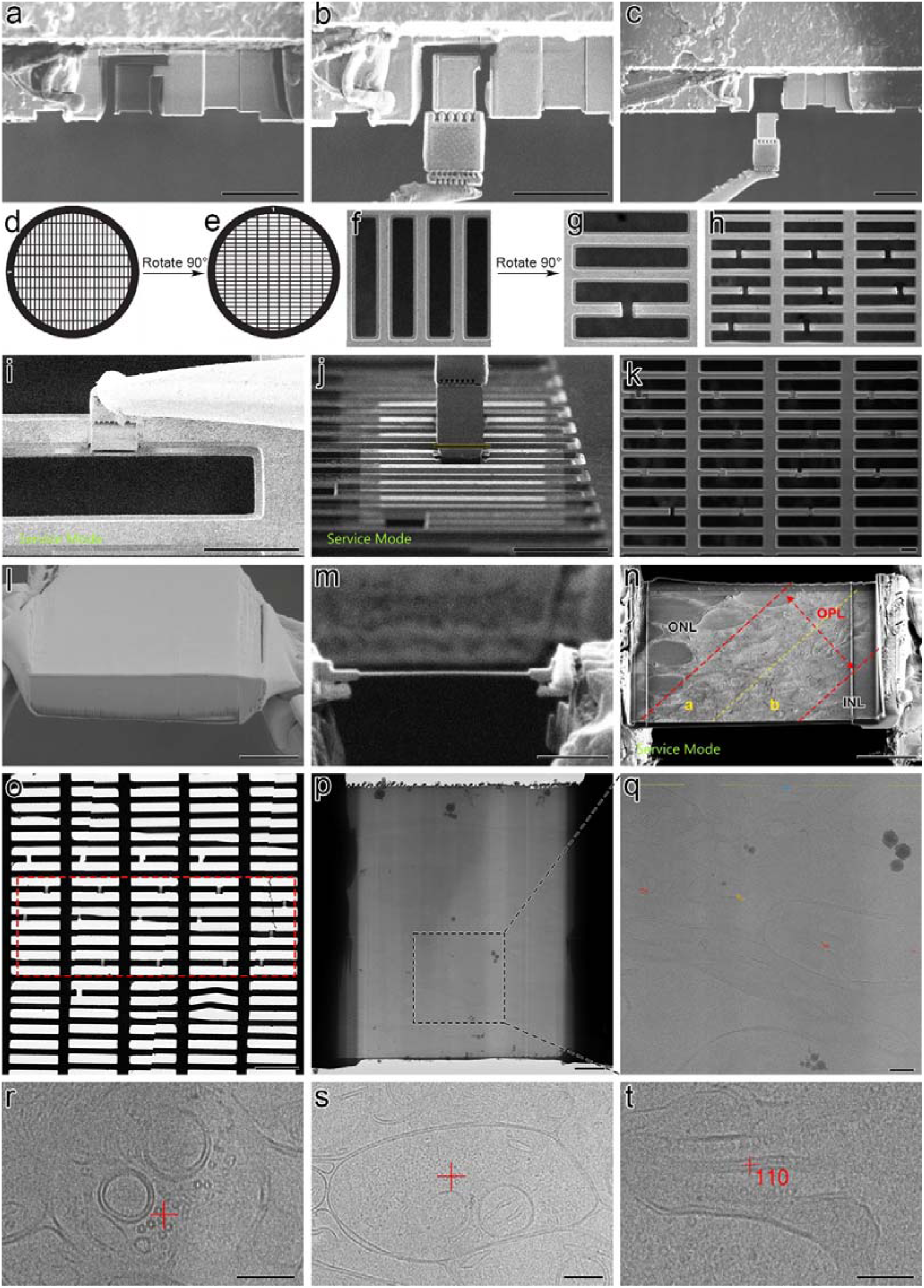
Lift-out, lamella preparation, and targeting selection. **a-c**, ROI extraction using an Easylift micromanipulator. The ROI block is separated from the carrier (**a**). The micromanipulator needle contacts the ROI block (**b**). The ROI block is lifted out from the carrier (**c**). Scale bars: 50 µm. **d-h**, TEM grid preparation. The grid is rotated 90° (**d, e**). A 20- µm-wide slot is milled into the grid bar (**f, g**). An overview of the prepared grid shows multiple milled slots (**h**). **i-k**, Sample attachment. The extracted block is positioned over the grid slot (**i**), segmented into ∼3–4- µm-thick sections (**j**), and each section is welded to the grid (**k**). Scale bars: 50 µm. **l-n**, Representative images at thinning stages. The unthinned extraction block (**l**), followed by side-view (**m**) and top-view (**n**) of the lamella. In the top-view, the OPL is demarcated by red dashed lines, with a yellow dashed line dividing it into areas a (ONL-proximal) and b (INL-adjacent). Area a contained round or oval profiles, presumably photoreceptor terminals, whereas area b displayed numerous irregular postsynaptic elements (Refer to Supplementary Fig 7a). Scale bars: 5 µm. **o-q**, Cryo-TEM imaging of the prepared lamella. Low-magnification (×125) survey image of the grid and the red dashed box indicates the position of the sample lamella (**o**). Medium-magnification (×8,700) image of an individual lamella (**p**) and enlarged view of the boxed region (**q**). Scale bars: o, 200 µm; p, 2 µm; q, 500 nm. **r**, Representative cryo-TEM image of microtubules in neuronal processes. Scale bars: 200 nm. **s**, Representative cryo-TEM image of neurofilaments within a horizontal cell axon. Scale bars: 200 nm. **t**, Representative cryo-TEM image of a synaptic ribbon in a rod spherule. Scale bars: 200 nm.

### Optimized cryo-FIB milling of the OPL

Initially, we directly placed extracted samples onto a 40- µm-wide grid mesh, but this frequently led to fractures during subsequent ion thinning. To overcome this, we rotated the grid by 90° before sample mounting and milled a 20- µm slot on the grid bar (Fig. 2d-h), as previously reported^25^. This slot width fully covered the OPL and enabled more stable thinning. The extracted sample was then cut into sections approximately 3–4 µm thick (Fig. 2i-l). We systematically optimized the ion beam milling protocol: the beam current was gradually reduced from coarse-milling to fine-thinning levels under high voltage, while the beam was kept nearly parallel to the carrier surface, so that the milling width was progressively narrowed to minimize stress and prevent bending. The resulting lamellae, ∼12–15 µm wide and ∼200 nm thick (side-view, Fig. 2m), were suitable for high-resolution cryo-ET. In the top view (Fig. 2n), the OPL is flanked by the ONL on the left and the INL on the right. Area a (near ONL) in the OPL contained round/oval profiles, presumably photoreceptor terminals, whereas area b (near INL) displayed irregular postsynaptic elements. Thus, this approach allowed precise OPL localization with clear visualization of presynaptic terminals near the ONL and postsynaptic elements on the opposite side, enabling rapid ROI identification.

### Cryo-ET data acquisition of synaptic ultrastructure in the OPL

Using a PLCT workflow, we acquired in situ cryo-ET data of synaptic ultrastructural elements within intact mouse retinal tissue. The grid was first surveyed at x125 (Fig. 2o), followed by imaging of individual lamellae at x8,700 (Fig. 2p, q). Minimal ice contamination and intact lamellar morphology enabled target selection, which was further refined based on retinal layer architecture and anatomical features within the OPL. We focused on microtubules and neurofilaments within postsynaptic horizontal cell processes (Fig. 2r, s), and presynaptic ribbons of rod terminals, which appeared as elongated rod-like structures (Fig. 2t).

### In situ subtomogram averaging (STA) of microtubules and neurofilaments

In cryo-ET images of the postsynaptic region, microtubules were observed within horizontal cell processes as bundled tubes (Fig. 2r). They frequently occurred in clustered arrangements, often near mitochondria, vesicles, and endoplasmic reticulum (Fig. 3a, b and Supplementary Video 1), consistent with a potential role in vesicle-associated intracellular transport^26^. Immunostaining confirmed that microtubules were enriched in horizontal cell processes (Supplementary Fig. 5a-a2). STA of extracted segments yielded a density map revealing a tubular architecture with 13 circumferentially arranged protofilaments (Fig. 3c and Supplementary Video 2) and a single solid central density along the longitudinal axis. The outer diameter was ∼25 nm, with an axial periodicity of 8 nm, matching the αβ-tubulin dimer repeat^26, 27^. The reconstruction reached 16.33 Å resolution (FSC 0.143, Fig. 3d), matching that of purified preparations and demonstrating that PLCT yields data of sufficient quality for reliable in situ STA in native tissue.

**Figure 3.**
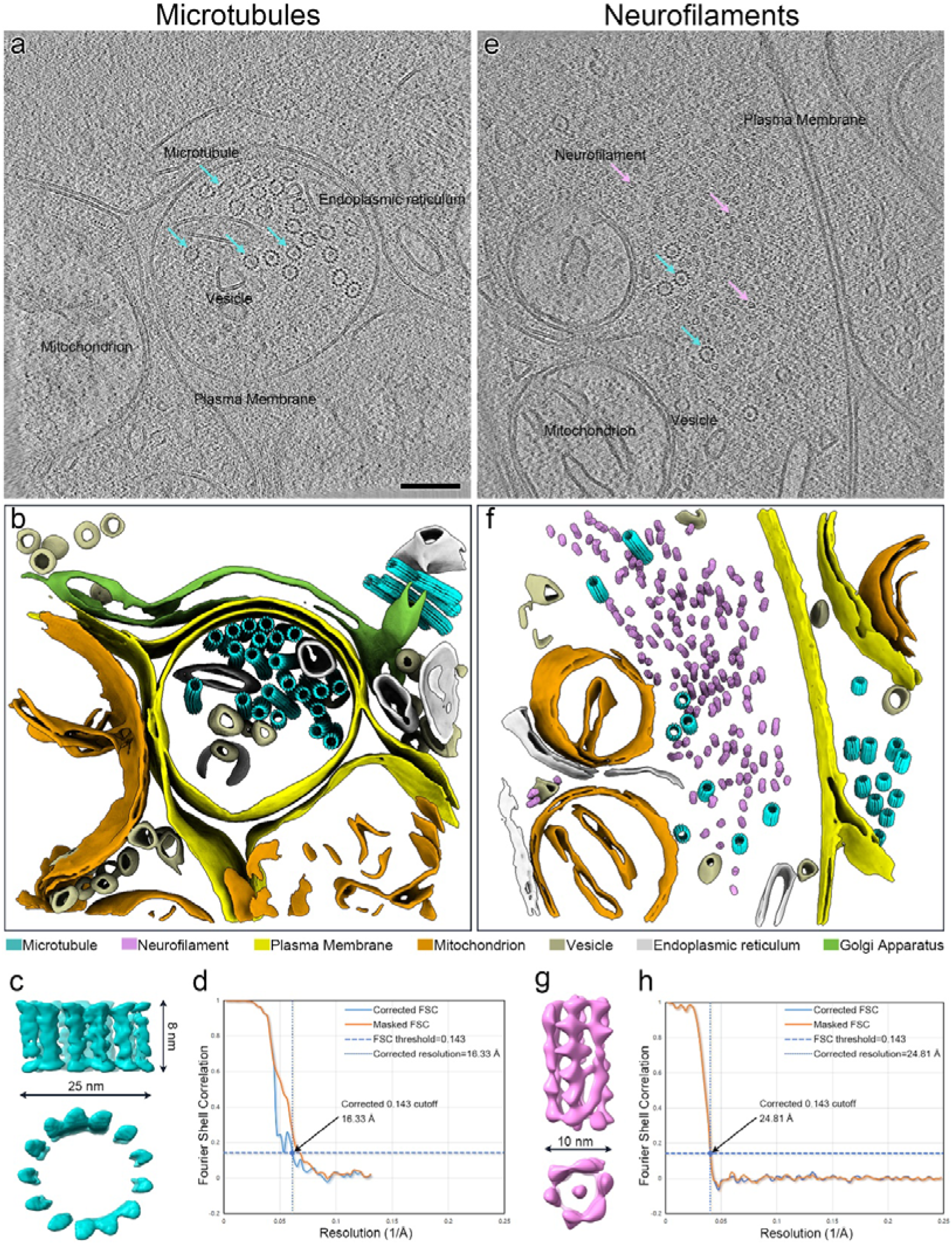
PLCT resolves native microtubules and neurofilaments in the mouse horizontal cell processes. **a-d**, Representative native microtubules. A representative reconstructed tomogram slice acquired from a PLCT section shows postsynaptic elements including horizontal cell processes having numerous microtubules, the cyan arrows indicate microtubules (**a**). The corresponding three-dimensional segmentation from **a**, highlighting the microtubules (cyan), plasma membranes (yellow), mitochondria (orange), vesicles (beige), endoplasmic reticula (grey), and Golgi apparatuses (green), thereby delineating the spatial organization of subcellular compartments within the tomographic volume (**b**). The surface rendering of the microtubule reconstruction viewed from two orthogonal orientations (top: side-view; bottom: top-view), revealing a tubular architecture with an outer diameter of approximately 25 nm and an axial periodicity of 8 nm (**c**). The Fourier shell correlation (FSC) curves for the microtubule subtomogram average, indicating a final resolution of 16.33 Å as determined by the 0.143 criterion (**d**). see also Supplementary Video 1, 2. Scale bars: 100 nm. **e-h**, Representative native neurofilaments. A representative reconstructed tomogram slice acquired from a PLCT section shows postsynaptic elements including horizontal cell processes having numerous neurofilaments, the magenta arrows indicate neurofilaments, while the cyan arrows indicate microtubules (**e**). The corresponding three-dimensional segmentation from **e**, highlighting the neurofilaments (magenta), microtubules (cyan), plasma membranes (yellow), mitochondria (orange), vesicles (beige), and endoplasmic reticula (grey), thereby delineating the spatial organization of subcellular compartments within the tomographic volume (**f**). The surface rendering of the neurofilament reconstruction viewed from two orthogonal orientations (top: side-view; bottom: top-view), exhibiting a tubular architecture with an outer diameter of approximately 10 nm (**g**). The FSC curves for the neurofilament subtomogram average, indicating a final resolution of 24.81 Å as determined by the 0.143 criterion (**h**). see also Supplementary Video 3, 4. Scale bars: 100 nm.

In addition to these bundled microtubules, thinner filamentous structures were also observed in the same horizontal cell processes (Fig. 2s), preliminarily identified as intermediate filament-like structures. These filaments typically formed bundled arrays in close proximity to microtubules and mitochondria (Fig. 3e, f and Supplementary Video 3). STA revealed a filamentous structure with an overall diameter of 10 nm, comprising six peripheral density strands around an elongated central density and continuous cavities between the strands. The reconstruction was resolved to 24.81 Å (Fig. 3g, h and Supplementary Video 4). These filaments share with vimentin a ∼10-nm tubular architecture and a central density, but are distinguished by six peripheral electron-dense strands, unlike vimentin’s five^28–30^. They were natively localized within horizontal cell processes, where neurofilaments have been previously reported to be expressed^31^. Our immunostaining further confirmed that neurofilaments are predominantly present in horizontal cells of the OPL (Supplementary Fig. 5b-b2). Taken together, the combination of structural characteristics, native localization, and immunoreactivity identifies these intermediate filaments as neurofilaments. Notably, microtubules were found in all regions that contained neurofilaments (Fig. 3e, f); however, some regions exhibited only microtubules and no neurofilaments (Fig. 3a, b and Supplementary Fig. 6).

### PLCT reveals native ribbon synapse ultrastructure

In the presynaptic region, a cryo-ET tomographic slice revealed a rod ribbon synapse within its native vitreous environment (Fig. 2t, 4a). The ribbon appeared as a segmented rod-like structure with a trilaminar organization: wide, high-contrast lateral flanks, a thin low-contrast central line, and numerous surrounding synaptic vesicles (Fig. 4a-c). Three-dimensional reconstruction further showed that the high-contrast flanks consist of many parallel arrays of small blocks, orderly arranged like mahjong tiles with both sides mutually parallel, and densely encircled by synaptic vesicles (Fig. 4b, c). This overall structure and arrangement highly comparable to those observed by conventional transmission electron microscopy (TEM) (Fig. 4d). Double immunofluorescence staining with antibodies against a terminal marker and a ribbon marker confirmed their identity as photoreceptor synaptic ribbons (Supplementary Fig. 5c-c2). Although these morphological and immunolabeling data are informative, in-depth STA of synaptic ribbons was not pursued here, as their irregular geometry and variable orientation demand extensive particle sampling. Undoubtedly, this analysis will be performed once sufficient samples are accumulated. Instead, our current focus is to establish PLCT as a reliable, feasible, and reproducible approach for native tissue structural studies.

**Figure 4.**
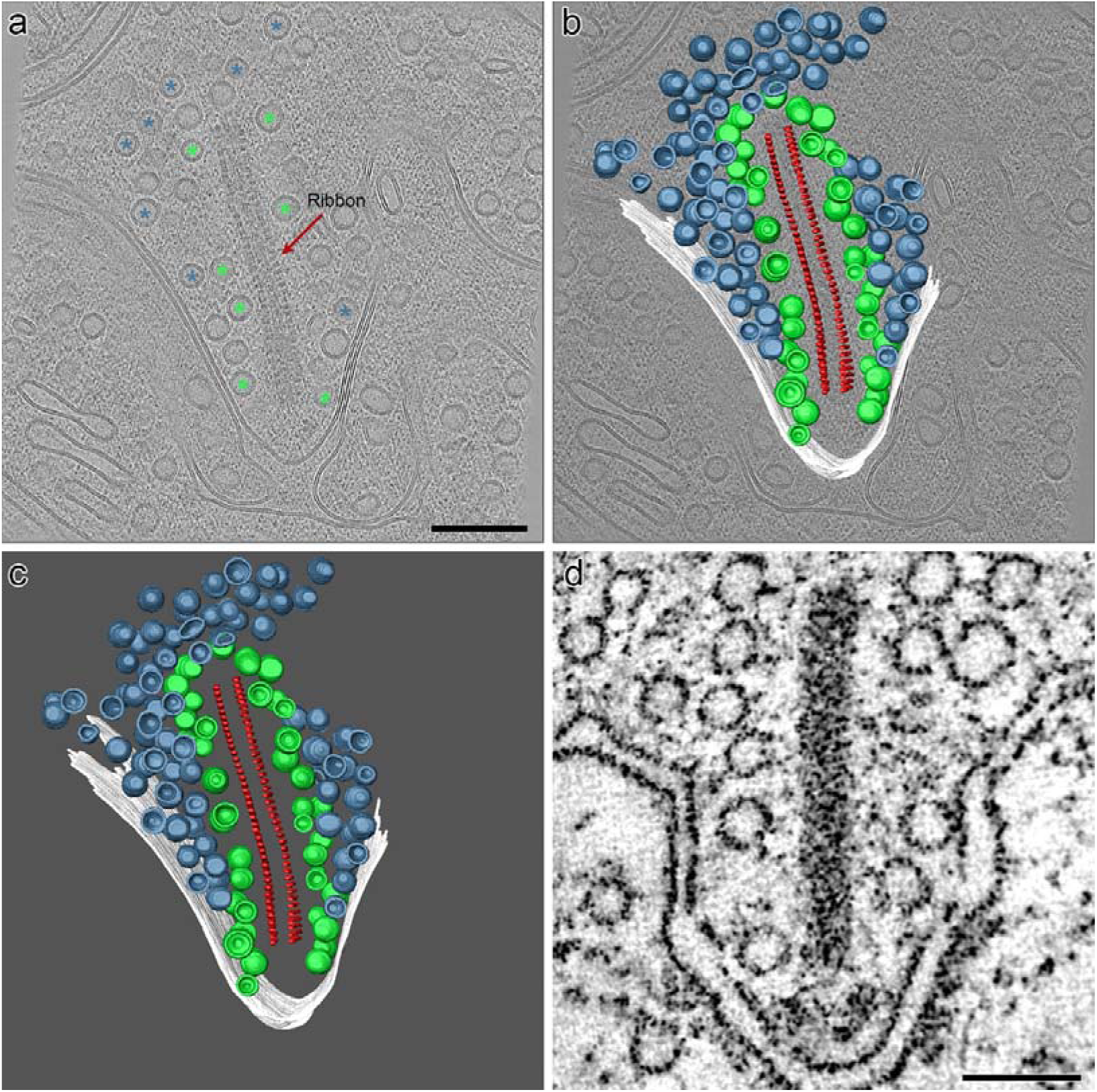
Representative native synaptic ribbon. **a-c**, A reconstructed tomogram slice from a PLCT section shows a presynaptic rod ribbon synapse, the red arrow indicates synaptic ribbon, green asterisks indicate ribbon-associated vesicles, and blue asterisks indicate reserve pool vesicles (**a**), with its three-dimensional segmentation overlaid in (**b**) and shown alone in (**c**). The ultrastructural components of the ribbon synapse: synaptic ribbons (red), vesicles (green and blue), and presynaptic membranes (white). Scale bars: a=b=c, 100 nm. **d**, A representative conventional TEM image displaying a rod synaptic ribbon. Scale bars: 100 nm.

## Discussion

To overcome the technical bottlenecks described above, we developed PLCT, a comprehensive workflow that integrates two highly feasible approaches: (1) dry-cutting HPF for robust vitrification of mouse retina, and (2) depth-controlled, layer-by-layer cryo-lamella preparation from large tissue blocks for in situ cryo-ET. For high success and efficiency, three critical practices are essential. First, retinal strips must be thinned to <100 µm (Fig. 5a, b), from the native ∼200 µm thickness, to lie flat on the substrate (Fig. 5c). Second, during transfer into the carrier, strips must be mounted without twisting or rotation (Fig. 5d); any deviation here causes cumulative positional errors later. Third, pre-trimming with a cryo-ultramicrotome must be perpendicular to the strip’s long axis (Fig. 5e), ensuring that every lifted-out section reliably contains the OPL (Fig. 5f, g).

**Figure 5.**
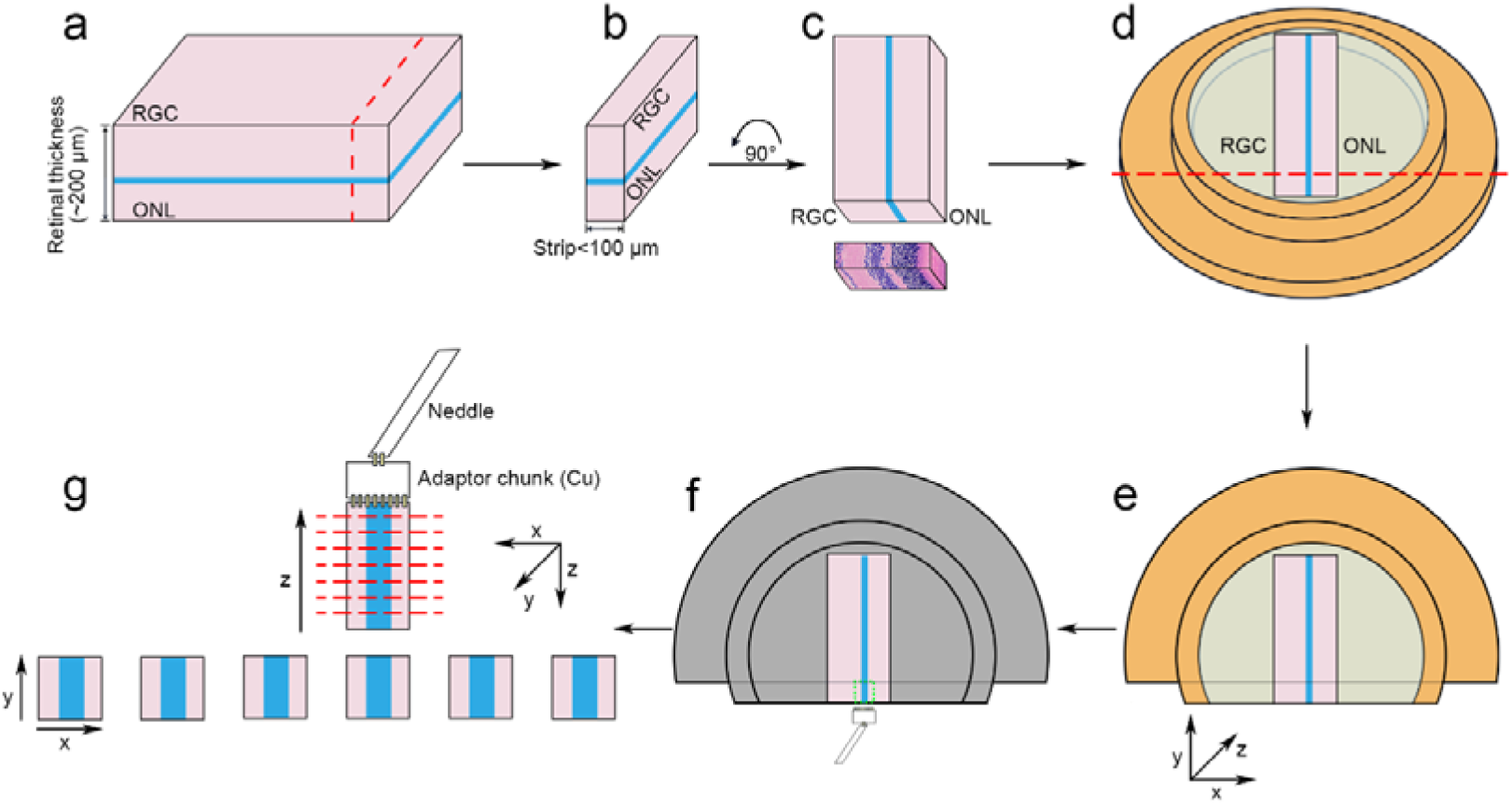
The main steps of PLCT. **a**, Retinal flat-mount section (200 µm thick). The OPL is highlighted in blue; red dashed lines indicate the cutting positions. **b**, Sectioned retinal strips (<100 µm thick). The OPL is oriented within the xy plane. **c**, The retinal strip is rotated 90° and laid flat, reorienting the OPL into the yz plane. **d**, The retinal strip mounted in the carrier. Red dashed lines indicate the positions for coarse pretrimming. **e**, Sample after fine trimming. The x-axis is defined perpendicular to the retinal strip, the y-axis along its longitudinal direction, and the z-axis along the carrier depth. **f**, Carrier setup in the cryo-FIB/SEM. The green dashed box indicates the region designated for lift-out. **g**, The lifted-out sample block. After lift-out, the original y-axis is redefined as the z-axis, and the original z-axis as the y-axis. The sample block is sectioned along the z-axis into multiple sections, containing the OPL in the xy plane.

### Dry-cutting HPF: a depth-resolved approach for multilayered tissue samples

Existing cryo-lamella and cryo-FIB workflows, developed for various tissues^32–34^ or optimized for cultured cells and thin sections^8, 12, 35^, are not directly applicable to the intact, multilayered retina. Chemical fixation and resin embedding introduce severe artifacts and limit resolution^36, 37^, while conventional HPF has only succeeded for the outermost photoreceptor discs^38^, leaving deeper synaptic regions, neuronal somata, and the ganglion cell layer entirely inaccessible. This failure stems from the increasing freezing depth and heterogeneous water/lipid composition of inner retinal layers, which impede uniform vitrification and subsequent FIB milling; consequently, no established HPF or plunge-freezing protocol currently enables depth-resolved in situ cryo-ET for these regions.

To bridge this methodological gap, we developed a modified dry-cutting HPF protocol that directly overcomes these barriers. This approach offers five critical advantages: (1) it eliminates vibratome sectioning, drastically reducing ex-vivo time to minimize hypoxic and mechanical damage^39, 40^; (2) it yields an optimal <100 µm thickness, avoiding the ice-crystal artifacts seen in thicker sections^36, 41^ while ensuring vitreous ice; (3) it omits agarose embedding, thereby bypassing infiltration artifacts such as uneven matrix distribution and altered contrast; (4) the combination of rapid processing and ideal thickness suppresses ice crystallization to negligible levels, safeguarding organellar integrity^42, 43^; and (5) the simple, low-cost workflow, requiring only a scalpel and dish, reduces operator variability and facilitates standardization. Collectively, this modified HPF ensures high-quality vitrification and faithful structural representation across all retinal depths, making robust in situ cryo-ET of inner retinal synapses and somata feasible for the first time and providing a decisive advantage for native-state imaging of neuronal and pathological features.

Beyond the retina, this method is not only applicable for precise localization of various cell layers or synaptic layers, but can also be used for accurate targeting of other tissues with similar layered structures or architectures, such as the cerebral cortex, cerebellar cortex, spinal cord white and gray matter, and renal cortex and medulla. Importantly, this approach is not limited to animal models; it can be adapted to human postmortem tissues including retinal specimens with appropriate consent and handling protocols.

### PLCT overcomes freezing and targeting barriers for depth**fll**resolved cryo-ET of layered specimens

Building on the modified HPF protocol described above, we further assembled a comprehensive pipeline, designated PLCT, that directly resolves two long-standing bottlenecks in tissue cryo-ET: (1) reliable vitrification of moderately thick specimens (100–300 µm), and (2) precise depth-targeted lamella preparation beyond the reach of conventional blind milling. PLCT integrates sequential steps—tissue dissection into <100 µm strips, modified HPF, cryo-ultramicrotome pre-trimming, morphological landmark identification, and xenon/argon plasma-based cryo-FIB milling—into a unified workflow. By reducing tissue thickness below 100 µm prior to HPF, we circumvent the non-uniform ice formation that typically plagues intact 200 µm retina. Subsequent cryo-ultramicrotome pre-trimming to ∼50 µm thickness bridges the gap between bulk tissue and electron-transparent lamellae while preserving native ultrastructure. Critically, the stepwise depth-navigation strategy, using morphological landmarks under the cryo-ultramicrotome followed by multimodal imaging in the cryo-FIB/SEM, enables unambiguous localization of the OPL at 90–100 µm depth, overcoming the “blind milling” limitation of conventional cryo-FIB that leaves intermediate targets inaccessible. Collectively, these innovations establish PLCT as a robust pipeline that transforms the retina—and, by extension, other layered tissues—from a challenging thick sample into a tractable specimen for depth-resolved in situ cryo-ET, thus holistically addressing the core HPF and cryo-FIB barriers that have long hindered inner tissue imaging.

### PLCT resolves native cytoskeletal and synaptic structures in mouse retina

Applying PLCT to the mouse retina enabled, for the first time, three-dimensional in situ reconstructions of both native cytoskeletal elements and synaptic structures within identified retinal components. Using STA on lamellae prepared from the OPL, we resolved microtubules to 16.33 Å and overall architecture closely matching those previously reported in other cell types and organisms^44–47^. Notably, this work also achieves the first in situ structural resolution of neurofilaments (24.81 Å) in a native tissue environment, a long-sought goal, as previous intermediate filament reconstructions (e.g., vimentin) were limited to in vitro or overexpressed systems^28–30^. The successful segmentation of these filaments within postsynaptic horizontal cell provides direct structural evidence for how they may organize synaptic machinery and maintain process architecture at the first visual synapse. Furthermore, we reconstructed the presynaptic photoreceptor ribbon synapse, including segmentation of the ribbon itself, opening a new window into the nanoscale organization of ribbon, vesicle tethering, docking, and release at this specialized active zone. Together, these results demonstrate that PLCT can resolve not only large membrane-bound organelles but also individual cytoskeletal filaments and synaptic protein complexes in their native tissue environment. The ability to unambiguously assign these structures to identified cell types (horizontal cells, photoreceptors) based on laminar position underscores the power of depth navigation for correlative structure–function studies in the retina.

### Strengths and inherent drawbacks of PLCT

Our PLCT workflow offers three key advantages for in situ cryo-ET of retinal tissue. First, it enables layer-by-layer milling at variable depths within large retinal samples, preserving the native layered architecture and providing access to both outer and inner structures, including synaptic terminals, neuronal cell bodies, and organelles. Second, the modified HPF protocol we developed specifically for mouse retina achieves consistent vitrification across the full tissue thickness, not merely the outermost photoreceptor discs. Unlike conventional HPF, which often fails to uniformly freeze deeper layers due to heterogeneous lipid and water content, our protocol yields ice quality sufficient for high-resolution cryo-ET throughout the retina. As a result, our workflow enables, for the first time, in situ cryo-ET visualization of inner retinal substructures, such as synaptic vesicles, cytoskeletal elements, mitochondrial membranes, and synaptic ribbons in their near-native state.

Despite its strengths, two limitations should be noted. First, dissecting retina into <100 µm strips may damage cut edges, so we removed approximately 25 µm from both the bottom and top surfaces of the tissue, leaving a ∼50-µm-thick slab for subsequent FIB milling. Second, ultramicrotome pre-trimming can introduce compression or chatter marks; this is mitigated by optimizing cutting parameters (e.g., ≤200 nm advance) and embedding in Ficoll, which we found superior to Dextran and bovine serum albumin (BSA). Nonetheless, PLCT provides a reproducible platform for high-fidelity cryo-EM of all retinal layers in health and disease.

### Future perspectives

PLCT relies on morphological landmarks for depth navigation and can also be applied to other multilayered tissues, extending its versatility across diverse sample types. Integration with cryo[Z]CLEM could further improve targeting precision, especially for rare or dynamically changing cellular events. Together, PLCT represents a major step forward, transforming thick[Z]tissue cryo[Z]ET from a formidable challenge into a practical and reproducible approach for investigating native macromolecular architectures within their authentic cellular context.

## Materials and methods

### Preparation of cryoprotectant and carrier

A 20% (w/v) Ficoll PM 70 (Sigma-Aldrich; F2878) solution was prepared by dissolving 200 mg in 0.1 M phosphate buffer (PB, pH 7.4) and adjusting the final volume to 1 mL. The solution was mixed thoroughly until completely dissolved, filtered through a 0.22 µm membrane filter, and used immediately to prevent degradation or contamination.

This study employed a customized cryo-carrier assembled with a specially designed auxiliary ring^48^. The carrier had an outer diameter of 3 mm, an inner diameter of 1.5 mm, and a depth of 100 µm. The carrier and sapphire plate were pre-cleaned by sonication in acetone for 10 min and air-dried in a dust-free environment. Before sample loading, the sapphire plate was inspected to confirm the absence of visible debris or residue.

### Animals

All animals used in this study were C57BL/6 mice (male, 8 weeks old, 20–22 g), purchased from Beijing Vital River Laboratory Animal Technology Co., Ltd. (Beijing, China). All animal procedures were approved by the Ethics Committee of Wenzhou Medical University and conducted in compliance with the ARVO Statement for the Use of Animals in Ophthalmic and Vision Research. Animals were housed under SPF conditions at 22 ± 2°C with a 12 h light/dark cycle and free access to food and water.

### Tissue acquisition and selection of the optimal approach for HPF

Mice were anesthetized by an intraperitoneal injection of 2,2,2-Tribromoethanol (250 mg/kg). A total of thirty-eight eyeballs were then enucleated. To optimize HPF preservation of retinal tissue, we systematically evaluated several preparation protocols (see also Supplementary Fig. 1). In the first approach, retinas were excised under oxygenated artificial cerebrospinal fluid (ACSF)^49^ and cut into flat-mounted sections (2 mm × 2 mm × 200 µm thick) for direct HPF. However, this method proved unsatisfactory because the 200 µm thickness exceeded the optimal freezing depth, resulting in conspicuous ice crystal formation (Supplementary Fig. 1a, a1). As an alternative, we embedded retinas in agarose and vibratome-sectioned them into <100 µm thick strips, but this was time-consuming and introduced infiltration artifacts (Supplementary Fig. 1b, b1). Ultimately, we adopted a refined protocol: retinas were rapidly cut into <100 µm thick strips (1.5 mm × 200 µm × 100 µm) on a dry plastic dish using a sharp scalpel, which minimized processing time and yielded excellent ultrastructure preservation with negligible ice crystals (Supplementary Fig. 1c, c1).

Detailed procedures for the ultimately adopted method are provided below. Under a dissecting microscope, after enucleation, the corneal limbus was perforated with a 1-mL syringe needle, and the anterior segment was excised along the perforation using scissors. The lens was then removed from the posterior eyecup with forceps. The eyecup was subsequently cut into appropriately sized rectangular section with a scalpel to fit the carrier, and the sclera and vitreous body were stripped away with forceps to isolate the retina. Immediately thereafter, cryoprotectant (20% Ficoll) was added, and the retina was trimmed into strips approximately <100 µm thick with a scalpel and promptly transferred onto the carrier for HPF (Fig. 1a-c). In this step, three critical technical considerations must be followed. First, retinal strips must be cut to <100 µm thickness. This ensures high-quality HPF sample preparation and provides the spatial foundation for subsequent localization. Second, when transferring retinal strips into the carrier, avoid any twisting or rotational misalignment. Even slight rotation at this stage will carry through the entire workflow, eventually causing major positioning errors and undermining the accuracy of target identification. Finally, HPF must be immediately performed using an HPF COMPACT 01 (Wohlwend Engineering, Switzerland). The auxiliary ring was assembled, the sapphire plate mounted on top ensuring no bubbles inside the carrier, followed by immediate vitrification. The vitrified sample was transferred from the HPF holder and stored under liquid nitrogen. To ensure sample quality, pressure, temperature, and freezing rate are strictly controlled (Fig. 1d). To prevent protein degradation, the interval from enucleation to retinal immersion in cryoprotectant must be strictly limited to <60 seconds. This preparation method achieves complete vitrification of retinal tissue without ice crystal formation, yielding high-quality frozen samples (Supplementary Fig. 1c, c1).

### Pre-trimming and coarse targeting of retina in the carrier

Pre-trimming were performed as described previously^48^, with minor modifications. Briefly, following HPF, the samples were transferred to a Leica EM UC7 cryo-ultramicrotome with the EM FC7 cryochamber (Leica Microsystem, Austria).

Under the eyepiece, the sapphire plate and auxiliary ring were removed (Fig. 1e). The retinal strip in the carrier was positioned perpendicular to the glass knife edge, and the carrier was fixed (Fig. 1f). The sample together with its carrier was first trimmed along the y-axis to approximately two-thirds of its original thickness using a glass knife (SPEED: 100 mm/s, FEED: 200 nm). Given that the retinal tissue was not compressed only when the strip thickness was maintained within 100 µm, the strip thickness must be verified before fine trimming. The knife edge was aligned with one edge of the retinal strip and set to zero (Supplementary Fig. 2a, a1, c, c1), then moved to the opposite edge to measure retinal thickness (Supplementary Fig. 2b, b1, d, d1). Based on mouse retinal thickness, 200 µm was adopted as the reference threshold: strips measuring within this limit were considered uncompressed and retained for subsequent experiments, whereas those exceeding 200 µm were discarded as compressed. The retinal strips selected through this method did not receive compression, which was verified in the subsequent lift-out step (Supplementary Fig. 3). Qualified strips were fine trimmed, excess material was sequentially removed from the sapphire plate face and the carrier bottom with a diamond knife (Diatome Trim 90, Switzerland; SPEED: 80 mm/s, FEED: 80 nm), yielding a final thickness of 30–50 µm (Fig. 1g-h). At this trimming stage, the cutting direction must remain orthogonal to the lengthwise axis of the retinal strips. Finally, the slice was first photographed, and then ImageJ software was used to approximately measure the horizontal position and orientation (ONL and GCL) of the retinal tissue within the carrier, allowing precise localization inside the cryo-FIB/SEM.

### Cryo-FIB milling

A cryo-FIB/SEM dual-beam microscope (Helios 5 Hydra CX system, Thermo Fisher Scientific, US) fitted with a complete cryo-system by FEl, including a rotatable cryo-stage cooled by an open nitrogen circuit served as the platform for cryo-FIB milling. Sample transfer was achieved by loading the carrier onto the shutter and introducing it into the chamber through a cryotransfer system maintained at −180 °C.

Because vitrified biological specimens are electrically insulating, a conductive platinum layer was first deposited by ion-beam sputtering to suppress charging-induced drift and distortion during SEM imaging and FIB milling. To further protect the lamella from ion-beam erosion and prevent curtaining artifacts during milling, a thick organometallic platinum layer was subsequently deposited via Gas Injection System (GIS) as a sacrificial protective coating. The final protocol followed a sandwich configuration: an initial 60 s Pt sputter coating, followed by 60 s GIS deposition, and a final 120 s Pt sputter coating.

The preliminary milling coordinates were determined from a vertical-view SEM image, based on the sample strip location previously measured in ImageJ and converted to absolute stage coordinates. Guided by the targeting coordinates, the top surface (xy-plane)/ lateral face (xz-plane) of the sample was coarsely milled (30 kV, 7.6 nA), followed by fine polishing with a reduced beam current (30 kV, 0.74 nA). The surface was then imaged to confirm the OPL position.

The ion beam incidence angle was optimized throughout the milling process. Coarse milling was conducted at 30 kV with beam currents ranging from 7.6 nA to 2 nA. During thinning, the ion beam was maintained nearly parallel to the carrier surface, with the acceleration voltage held at 30 kV and the current progressively reduced from 2 nA to 60 pA.

High-resolution imaging was performed using a through-the-lens secondary electron detector (TLD-SE) on a field-emission SEM. The imaging parameters were optimized to maximize surface detail while minimizing beam damage: 1 kV accelerating voltage for surface-sensitive contrast, 13 pA probe current to reduce specimen charging, and 50 ns pixel dwell time for rapid acquisition. Line integration (32/64 lines) improved signal-to-noise without scan-related artifacts. Images at 6144×4096 pixels provided ample spatial detail for quantitative morphometric analyses.

### Cryo-ET data acquisition and processing

Cryo-ET tilt series were collected on a 300-kV field emission Titan Krios microscope (Thermo Fisher Scientific) equipped with a Falcon 4i direct electron detector. After thinning, the grid was transferred into the TEM Autoloader for imaging. The grid was first surveyed at ×125 magnification, followed by imaging of individual lamella at ×8,700 magnification. Data acquisition was controlled using SerialEM. Tilt series were acquired at 64000x magnification at a physical pixel size of 1.912 Å/pixel using a dose-symmetric acquisition scheme. Prior to data acquisition, the sample pretilt was visually assessed and adjusted to approximately +10° or −10° to accommodate the geometry imposed by grid loading, with 3° angular increments used during acquisition. Two tilt schemes were employed: −65° to +55° (with a −10° pretilt) and −55° to +65° (with a +10° pretilt). Each tilt series comprised 40 projection images, with a total electron dose of 120 e^−^/Å² and a defocus range of −2 to −3 µm.

Data acquisition focused on photoreceptor rod ribbon synapses, which were identified primarily based on ultrastructural features established by conventional TEM (Supplementary Fig. 7). The synaptic ribbon itself, found mainly in rod spherules, served as the primary landmark due to its large rod-like morphology and high electron density. Postsynaptic horizontal cell processes were identified lateral to the ribbon, appearing electron-lucent with larger profiles and frequently containing vesicles. In contrast, bipolar cell dendrites were positioned basally or centrally beneath the ribbon, exhibiting small dendritic tips that generally lacked such vesicles.

### Data preprocessing, tilt-series alignment, and three-dimensional reconstruction

Raw tilt series were first preprocessed using Warp (https://pmc.ncbi.nlm.nih.gov/articles/PMC6858868/). Tilt-series alignment was subsequently performed using AreTomo2 (https://www.sciencedirect.com/science/article/pii/S2590152422000095?via%3Dihub), and the resulting alignment parameters were imported into Warp. Three-dimensional tomograms were reconstructed in Warp. Datasets with different binning factors were generated according to the requirements of downstream particle extraction and STA.

### Microtubule particle picking and STA

#### PyTom-based template matching of microtubules

The microtubule density map EMD-6351 from the Electron Microscopy Data Bank was used as the initial template for template matching by PyTom in the reconstructed bin4 tomograms (https://www.sciencedirect.com/science/article/abs/pii/S1047847711003492?via%3Di hub). Template matching was performed on 51 cryo-electron tomography datasets, yielding an initial set of 9,603 candidate microtubule particles.

The three-dimensional coordinates and Euler angles obtained from template matching were imported into Warp for subsequent subtomogram extraction. To generate two independent half-datasets, the tomograms were divided into odd and even groups while maintaining approximately equal particle numbers in the two subsets.

#### Bin4 and bin2 STA

Microtubule subtomograms were extracted from the bin4 datasets using Warp with a box size of 44 × 44 × 44 voxels and a pixel size of 7.648 Å/pixel. An initial three-dimensional reference was generated using the initial coordinates and Euler angles of all particles obtained from template matching. To minimize initial-reference bias, the reference was low-pass filtered to 60 Å. The subtomograms were further aligned in RELION 4 to ∼24 Å resolution using a criterion of 0.143 for the Fourier shell correlation (FSC) value (https://elifesciences.org/articles/83724). The particle coordinates and Euler angles obtained from alignment of the bin4 subtomograms were re-imported into Warp, and bin2 subtomograms were extracted at a pixel size of 3.824 Å/pixel. Subsequent alignment in RELION achieved an average resolution of 16.33 Å for the MT.

### Neurofilament particle picking and STA

#### Manually particle picking of neurofilaments

Previous studies have reported that neurofilaments exhibit an axial periodicity of approximately 21 nm and a diameter of approximately 10 nm^50, 51^ (https://pmc.ncbi.nlm.nih.gov/articles/PMC3516374/; https://rupress.org/jcb/article-abstract/94/3/592/19720/Visualization-of-a-21-nm-axial-periodicity-in?redirectedFrom=fulltext). Based on these structural features, neurofilaments were manually annotated in the bin4 tomograms. For each neurofilament, the start and end points were manually defined, and particle positions were oversampled at 7-nm intervals along the direction from the start point to the end point, thereby generating a continuous series of candidate neurofilament particles. The initial orientation of each sampled particle was determined from the vector pointing from the start point to the end point of the corresponding neurofilament and converted into the corresponding initial Euler angles.

To generate two independent half-datasets, complete neurofilaments were assigned to odd and even groups.

#### Initial STA of manually picked particles

The coordinates and initial Euler angles of the manually picked particles were imported into Warp. Alignment of the bin4 and bin2 subtomograms was subsequently performed using RELION. The resulting reconstruction was imported into M for additional optimization of particle poses and imaging parameters, yielding an average resolution of ∼30 Å for neurofilaments (https://www.nature.com/articles/s41592-020-01054-7).

#### PyTom-based template matching of neurofilaments and STA

The density map obtained from the manually picked particles was used as a template for template matching in PyTom. Candidate particles were selected based on their template-matching scores and spatial distributions. In total, approximately 3,396 neurofilament particles were identified from 12 cryo-electron tomography datasets.

The particle coordinates and Euler angles obtained from PyTom were imported into Warp. Bin4 and bin2 subtomograms were sequentially extracted and subjected to alignment in RELION, yielding a resolution of approximately 27 Å for neurofilaments. Finally, the reconstruction was imported into M for additional refinement, resulting in a final neurofilament density map with a resolution of 24.81 Å.

### Membrane segmentation and three-dimensional visualization

Membrane structures in the tomograms were automatically segmented using MemBrain-seg (https://doi.org/10.1101/2024.01.05.574336). Different membrane structures were further segmented using UCSF Chimera (https://onlinelibrary.wiley.com/doi/epdf/10.1002/jcc.20084). The final density maps were visualized using UCSF ChimeraX (https://onlinelibrary.wiley.com/doi/10.1002/pro.3943). The refined coordinates and Euler angles obtained from alignment were used to place the averaged microtubule and neurofilament density maps back into their corresponding positions and orientations (https://onlinelibrary.wiley.com/doi/10.1002/pro.4472). For better visualization, particles showing obvious deviations were manually removed.

### Three-dimensional reconstruction of synaptic ribbon

Tilt-series images underwent image alignment and assembly via IMOD software package (version 4.1.10). Subsequent volumetric reconstruction, segmentation, and surface rendering were carried out using Amira 6.8 (Thermo Fisher Scientific). Briefly, by isolating and highlighting specific structures from the tomograms, each structure can be assigned to a different material (label). Here, red represents the ribbon in the rod spherule, green represents the ribbon-associated synaptic vesicles (including readily releasable pool and reserve pool), blue represents cytoplasmic vesicles, and white represents the presynaptic membrane. After segmentation, the generated surface is created via Project Menu > Selection Labels > Generate Surface, applied and renamed with a .surf suffix, visualized by clicking Surface View, displayed in pure 3D form by toggling off the Ortho Slice overlay, and refined by adjusting material visibility and colors under Properties > Materials.

### Tissue preparation for Immunofluorescence

Tissue preparation was accomplished as described previously^52^. Briefly, four mouse eyes were enucleated, and the anterior segment and vitreous body were removed. The posterior eyecups were fixed in 4% paraformaldehyde in 0.1 M PB (pH 7.4) for 25–30 min at room temperature. Following fixation, eyecups were rinsed five times in 0.1 M phosphate-buffered saline (PBS, pH 7.4) for 10 min per wash, then cryoprotected through a graded sucrose series (10%, 20%, and 30% in PB, 60 min each) and stored overnight in fresh 30% sucrose at 4 °C. Tissue was embedded in OCT compound (Tissue Tek, Torrance, CA), snap-frozen in liquid nitrogen, and stored at −20 °C. Cryostat sections (25 µm, transverse plane) were cut and mounted onto glass slides.

### Immunofluorescence

Immunofluorescence staining was accomplished as described previously^52^. Cryosections were gradually brought to ambient temperature prior to staining, and a hydrophobic barrier was immediately traced around each retinal slice using a DARKO pen to confine reagent application. Tissue slices then underwent four successive rinses with PBS (10 min per wash). To suppress non-specific background, specimens were saturated with 5% normal donkey serum (NDS; Sigma, St. Louis, MO) in PBS for 90 min at room temperature (RT).

Primary antibody labeling was performed by exposing sections to a cocktail of 3% NDS, 1% bovine serum albumin (BSA; Sigma), and 0.3% Triton X-100 containing the respective primary antibodies. Following a 2-hour equilibration at RT, slides were transferred to 4°C for overnight incubation. The next morning, antibody solutions were recovered for potential reuse, and sections received six extensive PBS washes (10 min each) to remove unbound primary antibodies.

Fluorophore-conjugated secondary antibodies (405, 488, 594; Jackson ImmunoResearch, 1:500) diluted in 3% NDS with 0.3% Triton X-100 were then applied for 2 h at RT. After final PBS rinses, sections were mounted under coverslips using an anti-fade mounting medium containing DAPI for nuclear counterstaining and confocal microscopy.

### Confocal microscopy

Sections were imaged on a Zeiss LSM900 confocal microscope equipped with a 20×/0.8 NA objective and a 63×/1.4 NA oil-immersion objective. Low-magnification overviews of retinal architecture were acquired with the 20× objective. For OPL imaging, Z-stacks spanning 4–6 µm in depth were collected at 0.3- µm step intervals using the 63× oil-immersion objective. Image processing and adjustments were performed with Zen software (Zeiss).

### Antibody characterization

A rabbit monoclonal antibody targeting 68-kDa neurofilament light chain (Diagbio, db14746, clone DGR16339) was generated using a recombinant mouse 68-kDa neurofilament protein as the immunogen. The antibody was validated for western blotting, immunohistochemistry on paraffin-embedded sections, immunocytochemistry/immunofluorescence, and immunoprecipitation in human, mouse, and rat samples (Diagbio, Hangzhou, China).

A recombinant rabbit monoclonal antibody targeting beta III Tubulin (Abcam, ab221935) was generated using a synthetic peptide corresponding to the human beta III Tubulin protein as the immunogen, which is the carrier-free version of ab52623. This antibody specifically recognizes a ∼52 kDa protein (TUBB3; UniProt ID: Q13509) and has been widely used to label neuronal microtubules, assess neurite outgrowth, and distinguish axonal compartments in immunocytochemistry, immunohistochemistry, and multiplex imaging applications^53^.

A monoclonal antibody targeting CtBP2 (BD Biosciences, 612044) was generated using a recombinant mouse CtBP2 fragment spanning amino acids 361–445 as the immunogen. This antibody specifically recognizes a ∼48 kDa protein and serves as a marker for synaptic ribbons in the mouse retina^52^.

A mouse monoclonal antibody against calbindin-D-28K (Sigma-Aldrich, C9848, clone CB-955) was generated using purified bovine kidney calbindin-D-28K as the immunogen. Calbindin-D-28K, a highly conserved 28 kDa calcium-binding protein, serves as a marker for horizontal cells in the mouse retina^54^.

An anti-VGLUT1 antibody (Synaptic Systems, 135 307) was used to identify glutamatergic synaptic terminals^55^. This polyclonal antibody recognizes the C-terminus of rat VGLUT1 (SLC17A7) and has been validated by knockout.

### TEM

TEM was performed as previously described^56^. Briefly, three mouse retinas were isolated from eyecups and cut into small strips (1 mm × 2 mm), which were immediately fixed in a mixed fixative containing 2% PFA and 2% glutaraldehyde in 0.1 M PB (pH 7.4) for 12 h at room temperature. After washing with 0.1 M PBS, the strips were postfixed in 1% osmium tetroxide and stained with uranyl acetate. Subsequently, the tissue was dehydrated in a graded acetone series and embedded in Epon-812. To screen for regions of interest, 1-µm-thick semithin sections were stained with toluidine blue and examined under a light microscope. Finally, 70–90 nm-thick ultrathin sections collected on copper grids were examined using a HITACHI-7500 electron microscope (HITACHI, Japan). Images were acquired in the OPL.

### Reporting summary

Further information on research design is available in the Nature Portfolio Reporting Summary linked to this article.

### Data availability

Cryo-EM density maps have been deposited in the Electron Microscopy Data Bank (EMDB): EMD-82056 for the microtubule and EMD-82052 for the neurofilament. Corresponding raw cryo-EM micrographs have been deposited in the Electron Microscopy Public Image Archive (EMPIAR): EMPIAR-13777 (for EMD-82056) and EMPIAR-13778 (for EMD-82052). All data will be released upon publication.

## Supporting information

Official PDB Validation Report 1

Official PDB Validation Report 2

## Acknowledgements

We thank Youli Jian, Lei Sun, and Xixia Li for their assistance with HPF and cryo-ultramicrotome trimming at the Center for Biological Imaging, Institute of Biophysics, Chinese Academy of Sciences (CAS). We are also grateful to Tianjiao Wang, Shuoguo Li, and Yan Zeng for Cryo-FIB milling and Cryo-ET at the National Multimode Trans-Scale Biomedical Imaging Center, the Interdisciplinary Center for Biointelligence and Center for Biological Imaging, and the Core Facility for Protein Science, Institute of Biophysics, CAS.

## Author contributions

J.Z. conceptualized the work. F.L.W., X.L., J.Z., and L.H.Y. prepared samples and performed high-pressure freezing. J.Z., and F.L.W. contributed to data collection, analysis, and interpretation. F.L.W., X.Q.L., and B.L.R. performed immunocytochemistry. J.Z. and F.L.W. conducted TEM. F.L.W. drafted the Results and Methods sections. J.Z. wrote the final manuscript. J.Z., J.Q., and F.S. reviewed and edited the manuscript. All authors read and approved the final version.

## Funding

This work was supported by grants from National Key Research and Development Program of China (2022YFA1105503) and Start-up Foundation of Wenzhou Medical University (89219003).

## Competing interests

The authors declare no competing interests.

## Ethical approval

This study was conducted in accordance with the principles of the Declaration of Helsinki. Ethical approval was obtained from the Ethics Committee of Wenzhou Medical University, following ARVO guidelines for the use of animals in ophthalmic and vision research

---

**Supplementary Figure 1.**
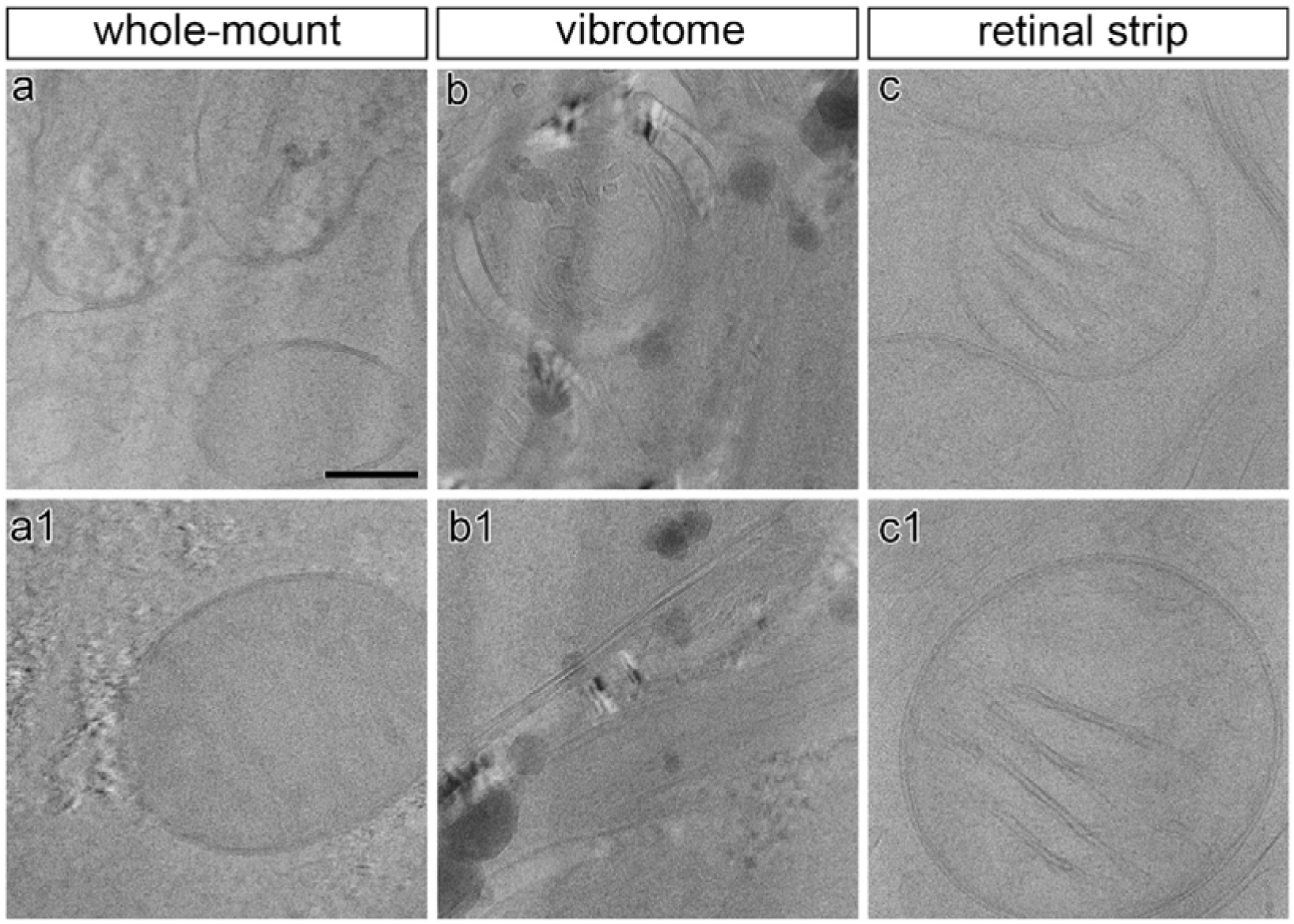
Evaluation of three HPF preparation methods. **a-c1**, Direct HPF of 200- µm-thick retinal flat-mounted sections results in suboptimal freezing with conspicuous ice crystal formation (**a, a1**). HPF of agarose-embedded retinas sectioned into 100- µm-thick strips using a vibratome still shows ice crystal artifacts despite reduced thickness (**b, b1**). Rapid cutting of 100- µm-thick retinal strips on a dry plastic dish yields satisfactory vitrification without ice crystals, enabling clear visualization of mitochondrial membranes (**c, c1**). Scale bars: 100 nm.

**Supplementary Figure 2.**
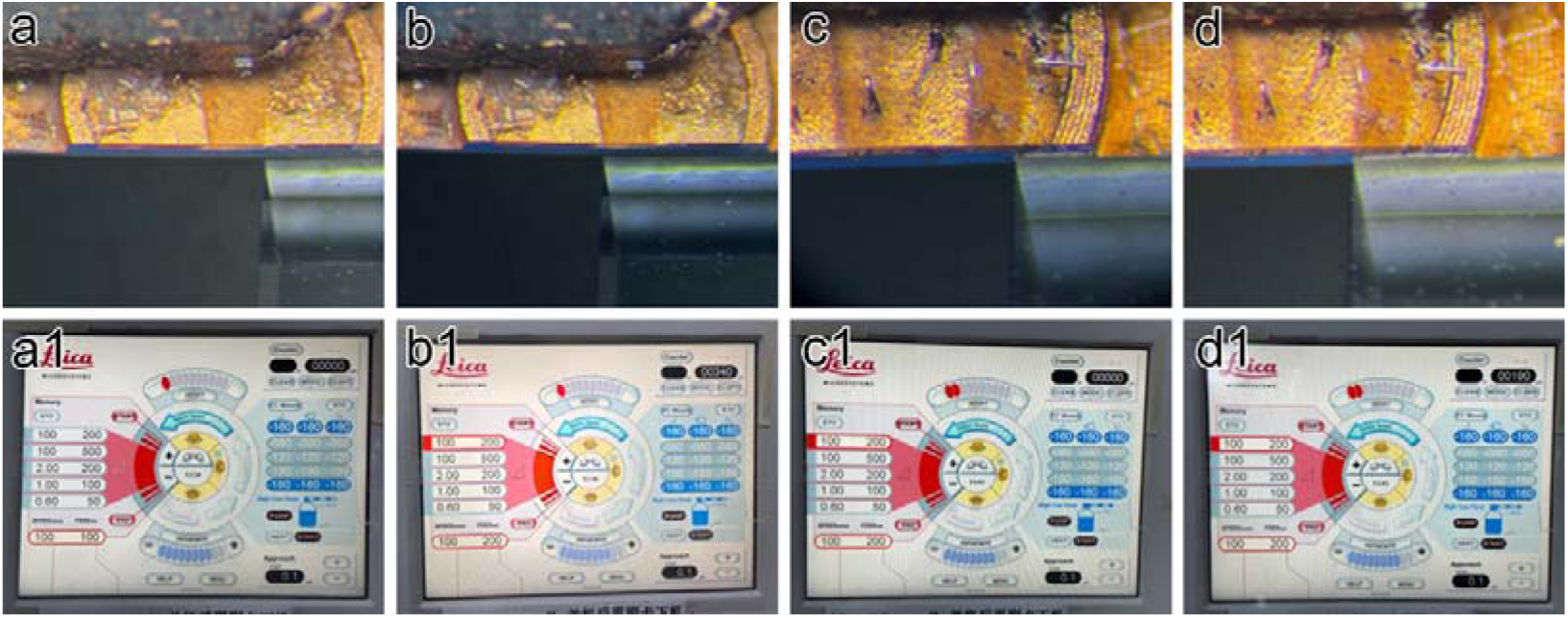
Strip thickness is inferred from retinal full-thickness measured by cryo ultramicrotome. **a-b1**, Representative measurement of a compressed retinal strip (strip thickness >100 µm because of retinal thickness >200 µm). The glass knife edge is aligned with one edge of the retinal strip and set to zero (**a, a1**). The knife edge is then moved to the opposite edge, and the measured displacement indicates retinal thickness (**b, b1**). **c-d1**, Representative measurement of an uncompressed retinal strip (strip thickness <100 µm because of retinal thickness <200 µm). The glass knife edge is aligned with one edge of the retinal strip and set to zero (**c, c1**). The knife edge is moved to the opposite edge, and the measured displacement indicates retinal thickness (**d, d1**).

**Supplementary Figure 3.**
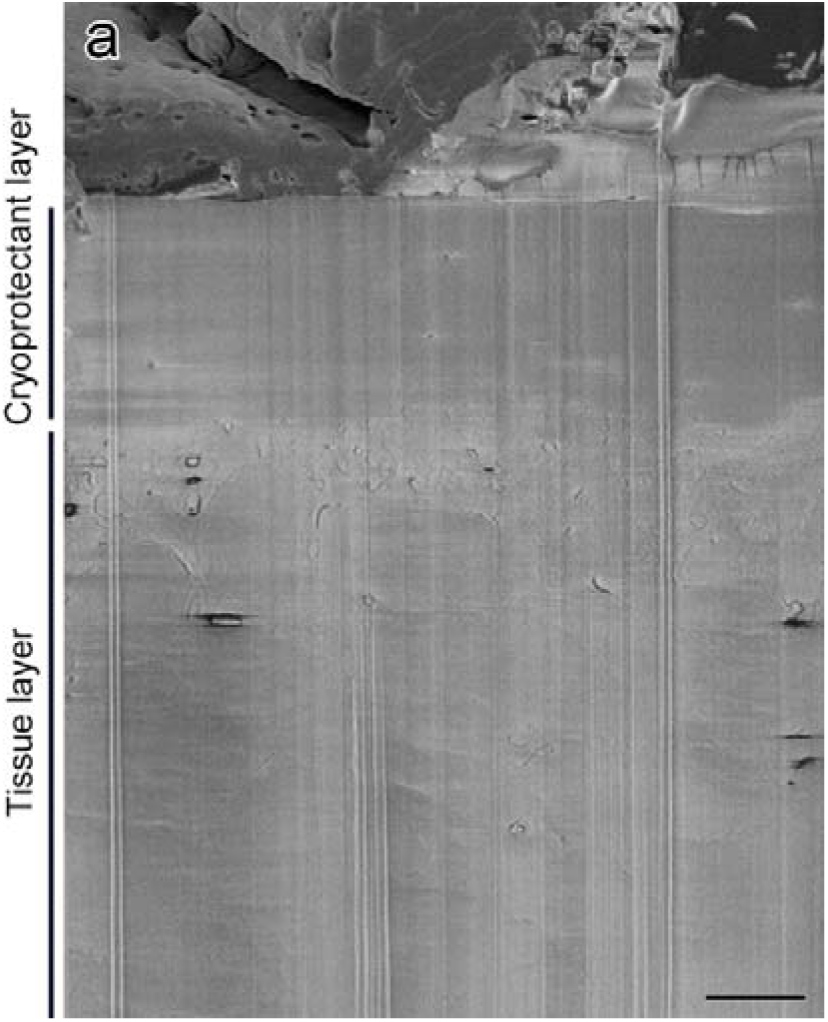
Validation of selected retinal stripes. **a,** A side-view cryo-FIB image after coarse milling and fine polishing clearly reveals the cryoprotectant and tissue layers, indicating no compression of the selected retinal strips. Scale bars: 2 µm.

**Supplementary Figure 4.**
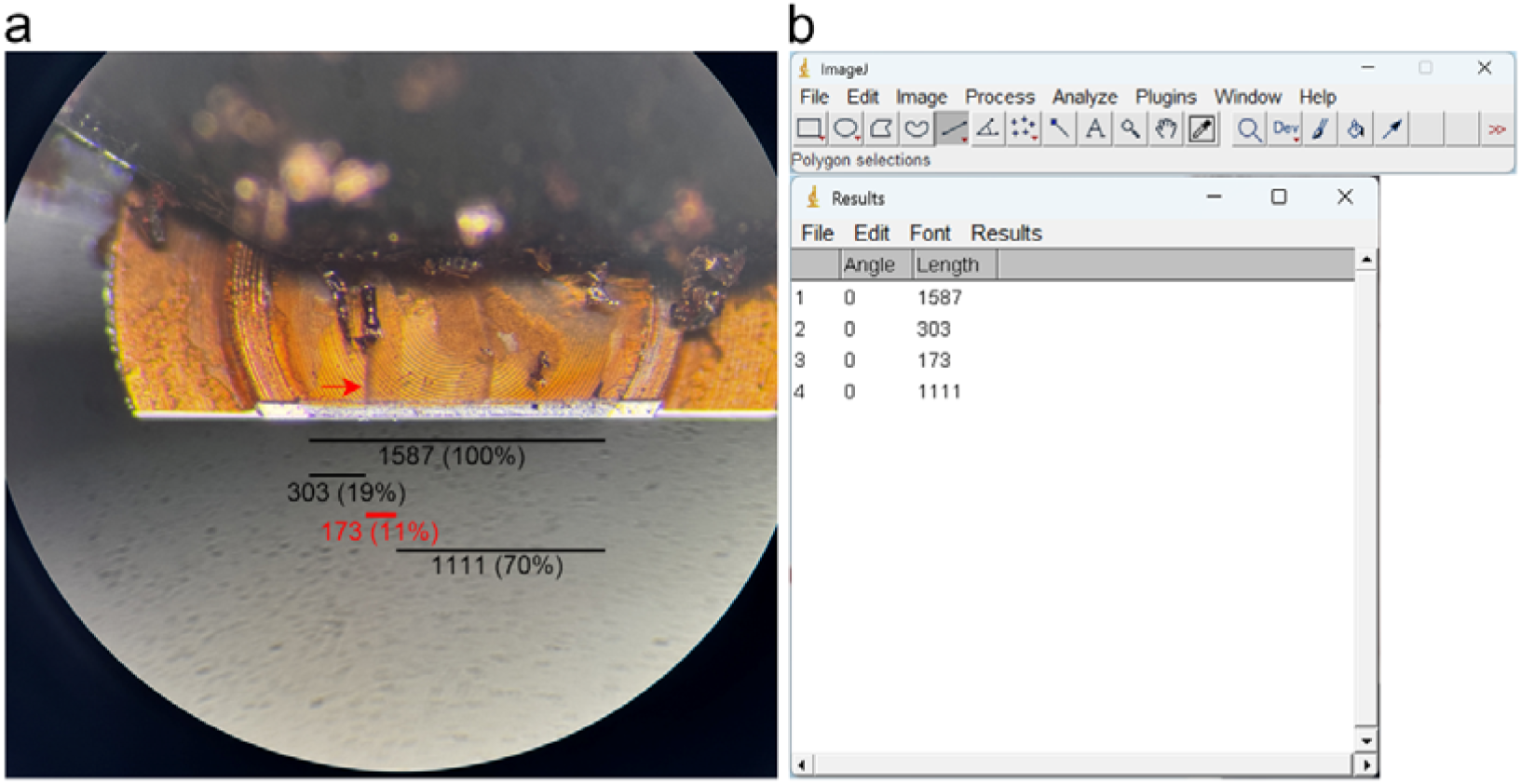
ImageJ-based percentage measurement for Hydra coordinate transfer. **a**, **b**, Bright-field images of pretrimmed carriers are imported into ImageJ. The relative position of the retinal strip (arrow) along the carrier x-axis is quantified as a percentage of the total carrier length (100%). In this example, the retina occupies 11% of the carrier, with 19% of the carrier length to its left and 70% to its right.

**Supplementary Figure 5.**
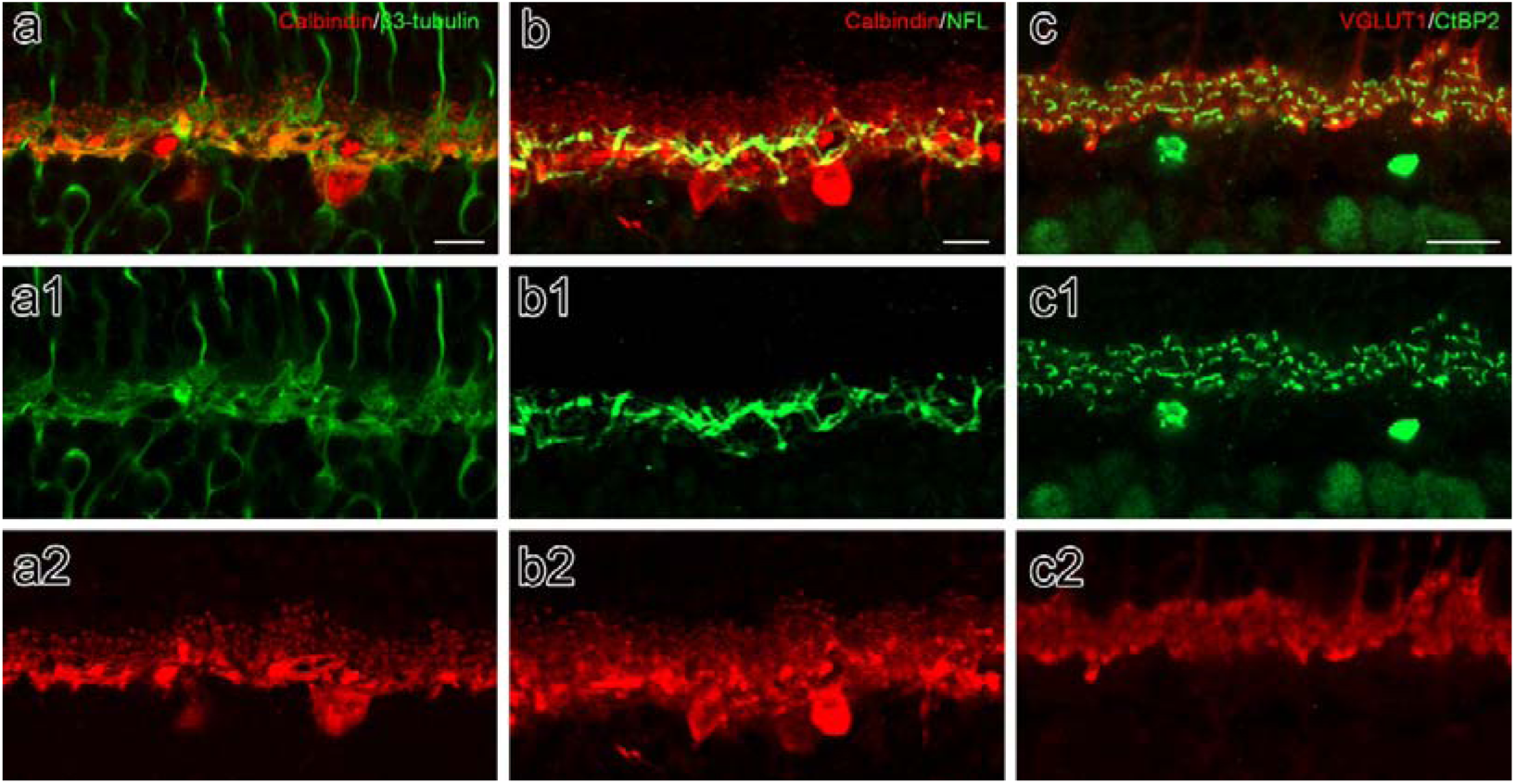
Immunostaining validation of targeting structures. **a-a2**, Representative double immunolabeling images showing ²3-tubulin (green, microtubule) and calbindin (red, horizontal cell) in the OPL. Microtubules are distributed within horizontal cell dendrites and axons. Scale bars: 10 µm. **b-b2,** Representative double immunolabeling images showing NFL (green, neurofilament) and calbindin (red, horizontal cell) in the OPL. Neurofilaments are confined to horizontal cell axons. Scale bars: 10 µm. **c-c2,** Representative double immunolabeling images showing CtBP2 (green, synaptic ribbon) and VGLUT1 (red, excitatory synaptic terminal) in the OPL. Synaptic ribbons are localized to VGLUT1-positive excitatory terminals (**c, c1**). Scale bars: 10 µm.

**Supplementary Figure 6.**
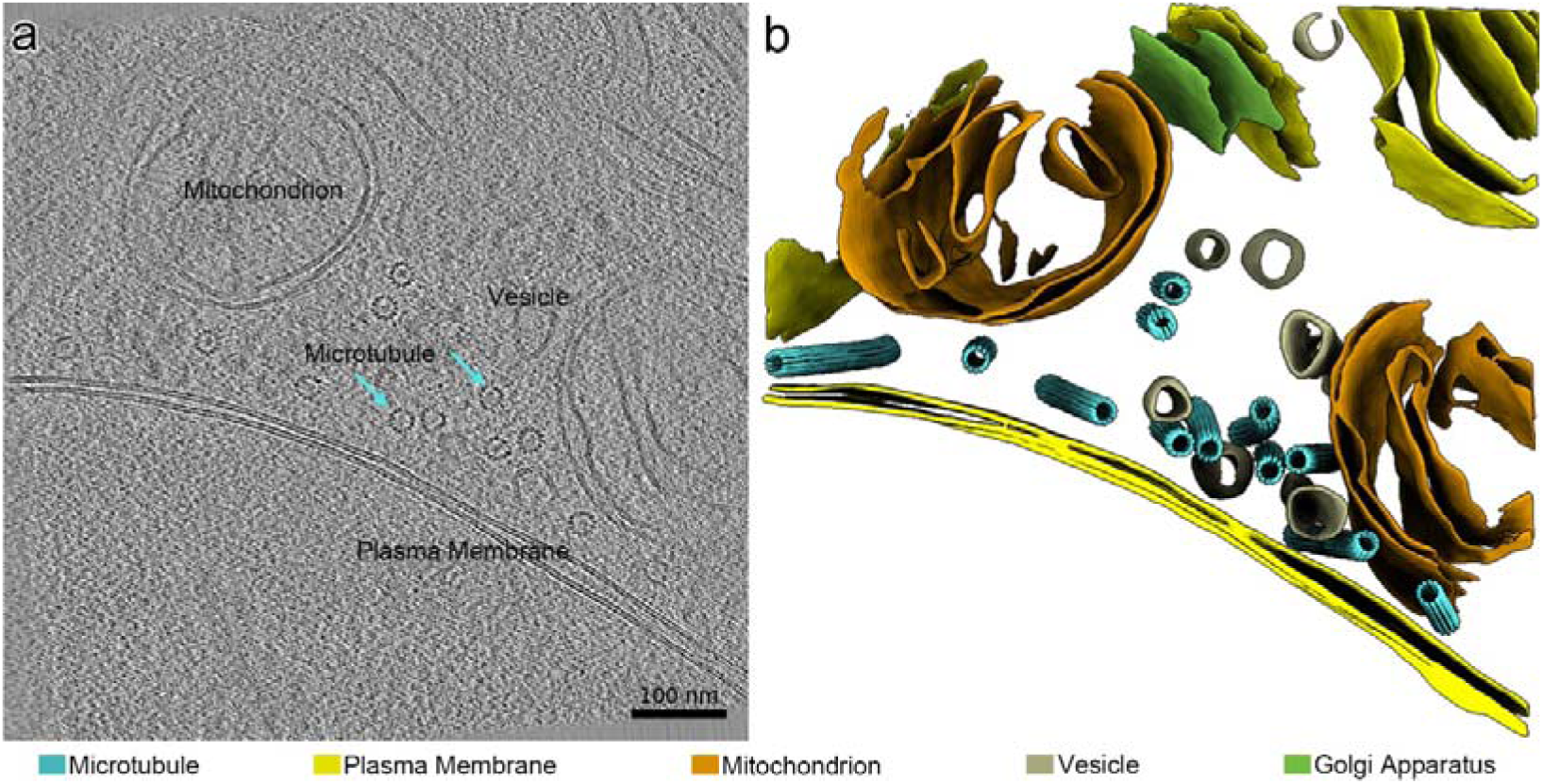
Representative native microtubules alone in the process. **a-b**, A representative tomogram slice (**a**) and its 3D segmentation (**b**) showing postsynaptic horizontal cell processes with microtubules (cyan) highlighted and no neurofilaments detected. Plasma membranes (yellow), mitochondria (orange), vesicles (beige), and Golgi apparatuses (green). Scale bars: 100 nm.

**Supplementary Figure 7.**
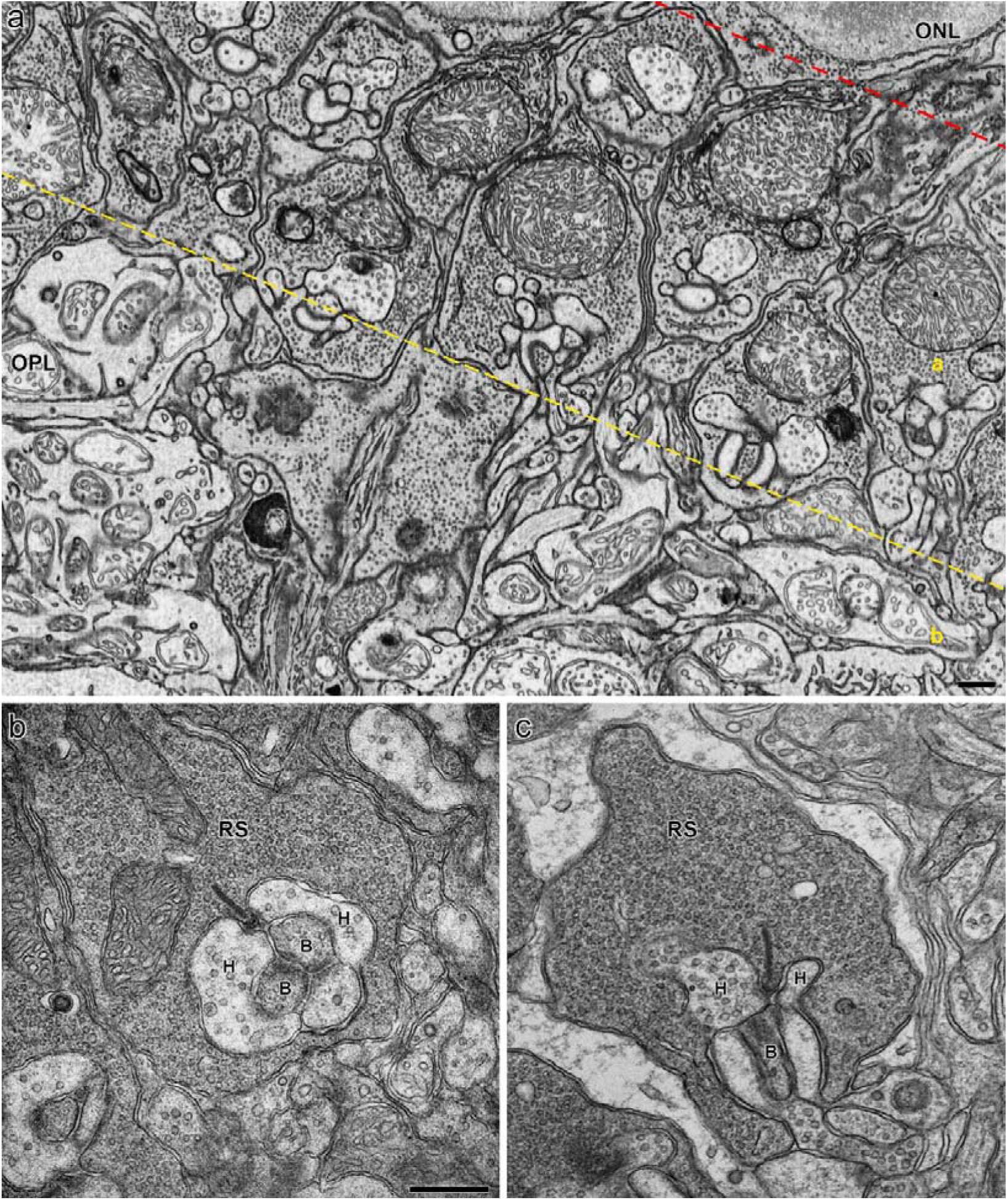
Representative conventional TEM images of the OPL and rod spherules in the mouse retina. **a,** A low-magnification TEM image shows the OPL area. A yellow dashed line divides it into a region (adjacent to the ONL) containing numerous photoreceptor terminals, and b region displaying irregular postsynaptic elements, comparable to that shown in Figure 2n. Scale bars: 200 nm. **b, c,** High-magnification TEM images reveal rod spherules. The presynaptic rod spherules contain a prominent synaptic ribbon, into which postsynaptic horizontal cell processes (H) and bipolar cell dendrites (B) invaginate. Horizontal cell processes, identified lateral to the ribbon, appear electron-lucent, exhibit larger profiles, and frequently contain vesicles. In contrast, bipolar cell dendrites are positioned basally or centrally beneath the ribbon and display small dendritic tips that generally lack such vesicles. Abbreviations: RS, rod spherule; H, Horizontal cells; B, Bipolar cells. Scale bars: b=c, 500 nm.

## Supplementary information

**Supplementary Video 1. Reconstructed horizontal cell process with microtubules.** This video shows a reconstructed horizontal cell process displaying multiple microtubules (cyan) surrounded by plasma membranes (yellow), mitochondria (orange), vesicles (beige), endoplasmic reticula (grey), and Golgi apparatuses (green).

**Supplementary Video 2. Reconstructed microtubule.** This video shows a reconstructed density map of a microtubule (cyan), displaying a tubular architecture with 13 circumferentially arranged protofilaments.

**Supplementary Video 3. Reconstructed horizontal cell process with neurofilaments and microtubules.** This video shows a reconstructed horizontal cell process displaying multiple neurofilaments (magenta) surrounded by microtubules (cyan), plasma membranes (yellow), mitochondria (orange), vesicles (beige), and endoplasmic reticula (grey).

**Supplementary Video 4. Reconstructed neurofilament.** This video shows a reconstructed density map of a neurofilament (magenta), displaying a tubular architecture comprising six peripheral subunits around a distinct central filament.

## Notes

### Competing Interest Statement

The authors have declared no competing interest.

